# Intensity-dependent effects of tDCS on motor learning are related to dopamine

**DOI:** 10.1101/2023.10.05.561136

**Authors:** Li-Ann Leow, Jiaqin Jiang, Samantha Bowers, Yuhan Zhang, Paul E Dux, Hannah L Filmer

**Author notes:** Corresponding author: Li-Ann Leow., Email address., Mailing address: School of Psychology, McElwain Building, The University of Queensland, St Lucia, Australia.

## Abstract

Non-invasive brain stimulation techniques, such as transcranial direct current stimulation (tDCS), are popular methods for inducing neuroplastic changes to alter cognition and behaviour. One challenge for the field is to optimise stimulation protocols to maximise benefits. For this to happen, we need a better understanding of *how* stimulation modulates cortical functioning/behaviour. To date, there is increasing evidence for a dose-response relationship between tDCS and brain excitability, however how this relates to behaviour is not well understood. Even less is known about the neurochemical mechanisms which may drive the dose-response relationship between stimulation intensities and behaviour. Here, we examine the effect of three different tDCS stimulation intensities (1mA, 2mA, 4mA anodal motor cortex tDCS) administered during the explicit learning of motor sequences. Further, to assess the role of dopamine in the dose-response relationship between tDCS intensities and behaviour, we examined how pharmacologically increasing dopamine availability, via 100mg of levodopa, modulated the effect of stimulation on learning. In the absence of levodopa, we found that 4mA tDCS improved and 1mA tDCS impaired acquisition of motor sequences relative to sham stimulation. Conversely, levodopa reversed the beneficial effect of 4mA tDCS. This effect of levodopa was no longer evident at the 48-hour follow-up, consistent with previous work characterising the persistence of neuroplastic changes in the motor cortex resulting from combining levodopa with tDCS. These results provide the first direct evidence for a role of dopamine in the intensity-dependent effects of tDCS on behaviour.

## 1. Introduction

Electrical brain stimulation approaches - such as transcranial direct current stimulation (tDCS) - can modulate cortical functioning and affect behaviour (Filmer, Dux, & Mattingley, 2014). An early example of stimulation influencing behaviour came from facilitatory effects of motor cortex tDCS on motor sequence learning (Nitsche et al., 2003). This classic finding for the field, however, does not always replicate in all populations (Ghasemian-Shirvan et al., 2023) and at times, stimulation has been shown to disrupt learning (Puri, Hinder, Krüger, & Summers, 2021). One possible reason for these inconsistencies is the use of varied stimulation protocols, driven by the fact we do not fully understand how stimulation modulates the brain. Indeed, it has recently been suggested that the some dosages (intensity) of stimulation applied to the motor cortex (0.7 - 1mA) may not be enough to produce reliable effects, and indeed intensities as high as 4mA may be required to achieve this (Hsu, Shereen, Cohen, & Parra, 2023; Shinde, Lerud, Munsch, Alsop, & Schlaug, 2021) .

The dose-dependence of tDCS effects on behaviour is incompletely understood. Whilst some studies suggest a linear relationship between stimulation intensity and effect of stimulation on behaviour (Filmer, Griffin, & Dux, 2019b; Shinde et al., 2021), other studies suggest a non-linear effect (Ehrhardt et al., 2022; Ehrhardt, Filmer, Wards, Mattingley, & Dux, 2021; Weller, Nitsche, & Plewnia, 2020). Yet other studies fail to show any behavioural difference between different stimulation intensities (Mitroi et al., 2020; Sommer & Plewnia, 2021), even when concurrently measured neurophysiological markers demonstrate dose-dependency (Ghasemian-Shirvan et al., 2023; Nikolin, Martin, Loo, & Boonstra, 2018). Whilst the finding that dose effects vary between targeted brain regions and associated cognitive processes is perhaps not surprising, more research is necessary to elucidate the neural mechanism giving rise to such effects of tDCS on behaviour.

One possible mechanism of action for tDCS dose-dependency on learning is the modulation of dopamine levels in the cortico-basal ganglia pathways. Indeed, there is evidence that tDCS may alter striatal activity (Chib, Yun, Takahashi, & Shimojo, 2013; Meyer et al., 2019) and midbrain dopamine release (Bunai et al., 2021; Fonteneau et al., 2018; Fukai et al., 2019; Tanaka et al., 2013), as pharmacological manipulations of dopamine strongly modulates the effects of non-invasive brain stimulation (Fresnoza, Paulus, Nitsche, & Kuo, 2014a; Kuo, Paulus, & Nitsche, 2008; Monte-Silva et al., 2009; Monte-Silva, Liebetanz, Grundey, Paulus, & Nitsche, 2010; Nitsche et al., 2006). For example, levodopa combined with 2mA motor cortex tDCS resulted in a 20-fold increase in the persistence of tDCS-induced neuroplasticity (Kuo et al., 2008). In addition, when tested against a fixed tDCS intensity, different doses of dopamine drug manipulations have non-linear effects on tDCS-induced neuroplasticity: only medium (and not small or large) doses of dopamine drugs have been found to alter the neuroplastic effects of tDCS (Fresnoza et al., 2014a; Fresnoza et al., 2014b; Monte-Silva et al., 2009; Monte-Silva et al., 2010). This suggests that neuroplastic effects of tDCS might depend on a sweet spot of brain dopamine levels.

If different intensities of tDCS elicit different dopamine responses in the brain, and if intensity-dependent tDCS effects on behaviour result from different dopamine responses, then dopamine drug manipulations on behaviour should interact with tDCS intensity – which has not been systematically assessed to date. Thus, here, in a pre-registered study, we explored whether different intensities of tDCS alters behaviour partly by soliciting different brain dopamine responses. We used a motor sequence learning paradigm as our model, given the recent debate about optimal dosages of stimulation for this task (Cuypers et al., 2013; Ghasemian-Shirvan et al., 2023; Shinde et al., 2021). First, we hypothesized that the effect of M1 tDCS on explicit motor sequence learning depends on tDCS intensity, anticipating that 4mA in particular would facilitate learning. Second, we hypothesized that the effect of M1 tDCS on explicit motor sequence learning depends on brain dopamine levels, and thus the administration of levodopa would modulate effects of M1 tDCS on sequence learning. We anticipated that this effect would depend on tDCS stimulation intensity. Third, given one previous finding showing associations between dopamine genotypes and explicit motor sequence learning in older adults (Schuck et al., 2013), we hypothesized that motor sequence learning may depend on brain dopamine levels. Thus, we predicted that in the absence of stimulation (sham stimulation, see below), increasing dopamine availability would modulate motor sequence learning.

## 2. Materials and Method

### 2.1. Design

In this pre-registered study (https://osf.io/jegsr), we employed a factorial design, with between-subjects variables of dose (sham, 1mA, 2mA, 4mA) and drug (levodopa, placebo), and within-subjects variables assessing repeating versus random sequences, as well as the different study blocks. Briefly, all participants were asked to execute a repeating or a random sequence across baseline, training, test, and retention blocks (see Figure 1). Participants were randomly assigned to one of the eight conditions as shown in Table 1. In all sessions participants completed a 5-element sequence learning task, described below and outlined in Figure 1 & 2.

**Table 1:** Participants were randomly allocated to one of the 8 study conditions, where they received a placebo or levodopa tablet, and received either sham, 1mA, 2mA, or 4mA tDCS during the training block of the task.

|  |  | Stimulation |  |  |  |
| --- | --- | --- | --- | --- | --- |
|  |  | Sham | 1mA | 2mA | 4mA |
| Drug | Placebo | N = 17 | N = 17 | N = 17 | N = 16 |
|  | Levodopa | N = 17 | N = 16 | N = 19 | N = 18 |

**Figure 1.**
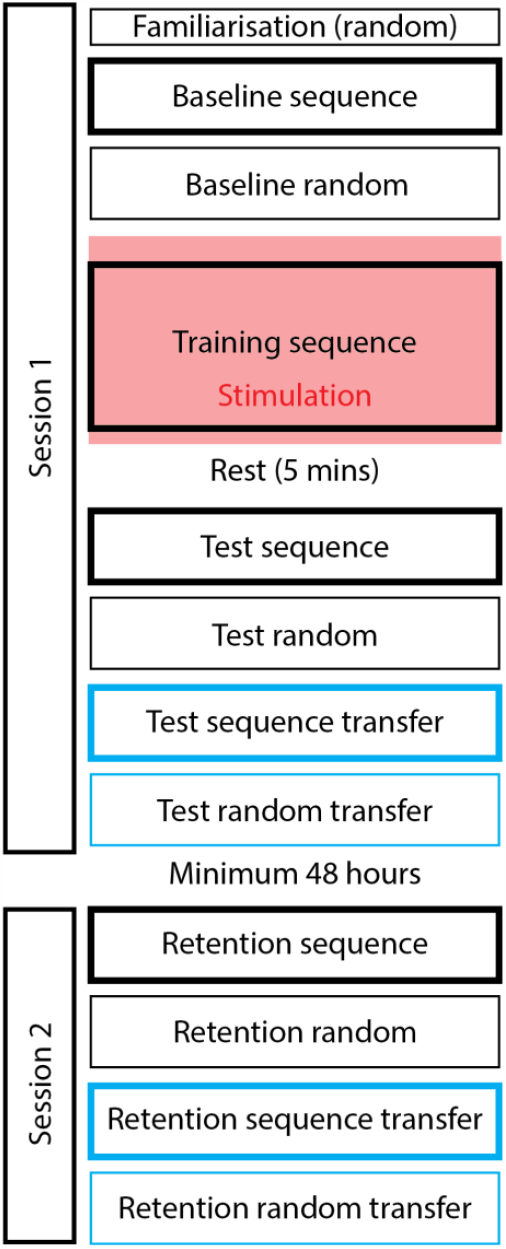
Task structure for session 1 and session 2. On session 1, participants executed a 5-element repeating sequence, or a 5-element random sequence at baseline, training, and test, using their left hand. In transfer blocks, they executed sequences with their untrained right hand. Stimulation occurred during training, After a minimum 48 hour interval, participants returned for session 2, where retention was first assessed for the trained hand, and then for the untrained hand (transfer blocks)

**Figure 2.**
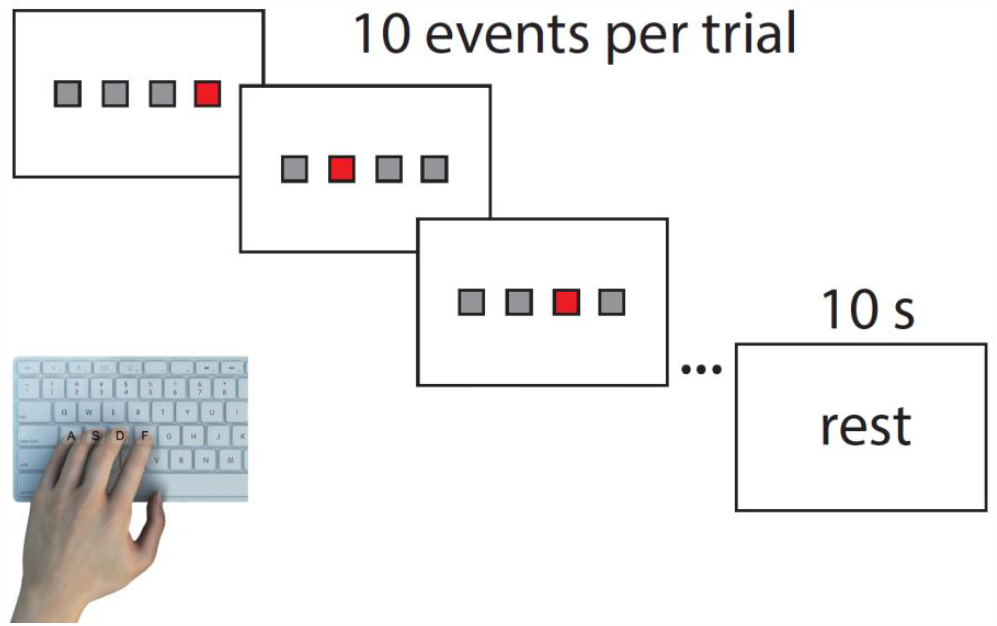
During the sequence learning task, participants saw 4 grey boxes on-screen which corresponded to the four finger positions on the keys ASDF. During sequence trials, participants executed a 5-element sequence (FSDFA) by responding as quickly and as accurately as possible to the red target stimulus, which appeared on-screen according to the sequence order. In random trials, the red target stimulus appeared in random order. During the training block, participants executed 28 bins of 10 trials: each bin of 10 trials was followed by a 10 second rest break.

### 2.2. Participants

Participants were right-handed and aged between 18 and 35 years old (mean age = 20 years, SD = 4 years), with no known neurological or psychiatric conditions and no contraindications to brain stimulation or Levodopa. In addition, as sequence learning in experts is differentially altered by tDCS compared to non-experts (Furuya, Klaus, Nitsche, Paulus, & Altenmüller, 2014), participants were required to have fewer than 13 years of musical training, and currently engaging in no more than 20 hours of musical training or gaming per week. Participants were pseudorandomly allocated to the following groups (see Table 1): levodopa sham tDCS (n=17), levodopa 1mA (n=16), levodopa 2mA (n=19), levodopa 4mA (n=18), placebo sham tDCS (n=17), placebo 1mA (n=17), placebo 2mA (n=17), placebo 4mA (n=16). The study was approved by the Human Research Ethics Committee at The University of Queensland and conformed to the Declaration of Helsinki. All participants provided written informed consent.

We adopted an adaptive Bayesian sampling plan. Our stopping rule stipulated that we were to recruit participants until a BF_10_ > 3 or BF_01_ > 3 was established for the critical hypothesis tests, providing moderate evidence for the alternative or null hypothesis for an effect of dose, or until we collected 30 complete datasets for each condition (total N=240), whichever was sooner. Specifically, we tested for evidence ***of*** a dose-dependent effect of tDCS on explicit motor learning, by examining if a Bayesian t-test on sequence-specific learning shows moderate evidence (Bayes Factor >3) **for a difference between any two of the three dosages**, or if there was moderate evidence (BF_incl_ >3) for including the main effect of dose in the Bayesian ANOVA. We tested for evidence ***against*** a dose-dependent effect of tDCS on explicit motor sequence learning by examining if Bayesian t-tests on sequence-specific learning showed moderate evidence (BF_01_>3) against differences between all of the three dosages, or if there was moderate evidence (BF_excl_>3) for excluding the main effect of dose in the Bayesian ANOVA. BF values were tested once a minimum of 15 participants were tested for each condition. We ended up with a sample size of between n=16 to n=19 per condition (total: n=138). As per our pre-registration, we discarded 3 datasets from the analyses due to poor task performance (accuracy less than 60% across several phases of the experiment).

### 2.3. Drug and stimulation manipulations

In session 1, participants first completed blood pressure and mood assessments, and then received either placebo (vitamin) or levodopa (Madopar 125: 100 mg levodopa and 25 mg benserazide hydrochloride), crushed and dispersed in orange juice. An experimenter uninvolved in data collection prepared the solution. This procedure was sufficient to achieve double blinding in previous work (Leow, Bernheine, Carroll, Dux, & Filmer, 2023a; Leow et al., 2023b). tDCS electrodes were then set up, before participants completed a second blood pressure and mood rating assessment, and a brief questionnaire assessing chronotype (the Morningness– Eveningness Questionnaire, DMEQ), which has recently been shown to modulate responsivity to brain stimulation and performance on motor sequence learning tasks (Salehinejad et al., 2021).

Participants then started the task component of the session (see Figure 1). They completed the familiarisation and the baseline blocks of the sequence learning task. Stimulation commenced – approximately 30 minutes after drug ingestion – within the window of peak plasma availability (Contin & Martinelli, 2010). Upon stimulation commencement, participants completed the training block, which assessed acquisition of the sequence. Stimulation lasted for 10 minutes, which coincided with the completion of the training block. After a 5-minute rest break during which electrodes were removed, participants completed an end-of-acquisition block. Blood pressure and mood rating assessment were again completed (approximately 1 hour after drug ingestion). After an overnight interval (minimum 48 hours, sufficient for effects of the drug and the stimulation manipulations to dissipate), participants returned to the lab for a no-drug, no-stimulation follow-up session.

For the 1mA, 2mA, 4mA conditions, stimulation lasted for 11 minutes (30 second ramp up, 10 minutes constant, and 30 seconds ramp down). For the sham condition, the parameters were the same but the stimulation lasted for 1 min 15 s (stimulation intensity was evenly split between 1, 2, and 4mA) followed by regular small test ‘pulses’ to maintain some sensation and allow impedance values to be calculated and displayed. Whether stimulation was active or sham was double blinded using stimulation codes provided by a team member who did not participate in data collection. Stimulation intensity was only single blind, due to limitations of the stimulator. Stimulation was delivered via a NeuroConn stimulator with two 5 × 5 cm electrodes. The target electrode (5cm x 5cm) was placed over the EEG 10-20 location C4 (Jasper, 1958), and the reference electrode (5cm x 7cm) was placed over the contralateral supraorbital region (the area above the left brow ridge; see Figure 3). Both electrodes were encased in saline-soaked sponge pads with a layer of highly conductive saline gel (SignaGel) (Kane, Sherwood, & Weisend, 2015)

**Figure 3.**
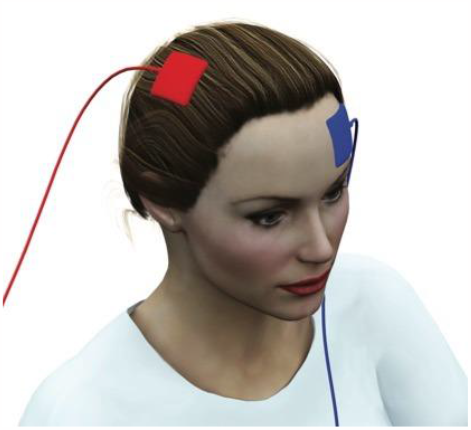
tDCS montage, with the 5cm x 5cm anode placed over the C4, targeting the right primary motor cortex, and the reference electorde (5cm x 7cm) placed over the left supraorbital area.

**Figure 4.**
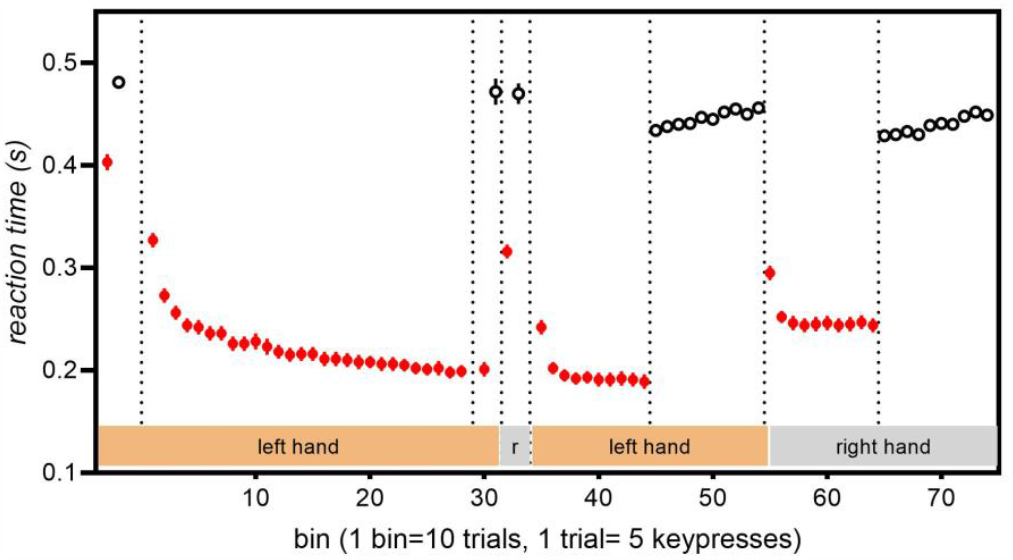
Reaction times for correct keypresses throughout all study phases. Red symbols = sequence trials, clear symbols = random trials. Lower values indicate greater performance improvement.

### 2.4. Task

Participants were first asked to place their little, ring, middle, and index fingers of their left hand on the keys ASDF. A row of four boxes were presented on screen, three grey and one red target stimulus, each corresponding spatially to the finger positions. The participants’ task was to respond as quickly as possible to the red target stimulus by pressing the appropriate key: in each trial, participants executed 5 keypresses according to the spatial location of the red target. There were two types of trials: sequence trials and random trials. In each sequence trial, the grey boxes turned red 5 times according to the 5-element tap sequence for the sequence blocks [FSDFA]. The sequence was the same throughout all trials for all participants. In each random trial, the grey boxes turned red 5 times in random order. At the start of each trial, brief prompts denoted trial commencement: “ready? (300ms) “start!” (300ms duration). No performance feedback was presented throughout the entire study.

First, participants were given the following on-screen instructions, which were read out loud by the experimenter “In this study, your task is to respond using the below keys when the corresponding box is highlighted red, using your LEFT hand. Press any key to start the familiarization block”. Then, participants completed a familiarization block (3 trials) with a random sequence to familiarise them with the task. Upon completing the familiarization block, participants were given the following on-screen instructions, which were read by the experimenter to ensure participant comprehension: “In some trials, the cues will appear in a fixed sequence. Respond to the cues as QUICKLY and ACCURATELY as possible.”

After familiarization, participants completed the baseline sequence block (10 trials, i.e., 10 executions of the 5-element sequence), followed by a baseline random block (10 trials). Stimulation commenced and was maintained for 2 minutes, before the start of the training block, which assessed acquisition of the repeating sequence. The training block consisted of 280 sequence trials (or 28 bins), during which a timed 10 second rest break was presented after every 10 trials. The training block was followed by a 5-minute rest break, during which the stimulation ended and the electrodes were removed. Participants then completed the end of acquisition block (i.e., the test block), which consisted of 1 sequence bin and 1 random bin (10 trials for each bin, or 1 bin) with the training hand, and 1 sequence bin and 1 random bin for the untrained right hand. This concluded the first session (Session 1).

After an interval of minimum 48 hours (sufficient time for effects of tDCS and levodopa to wash out), participants returned to the lab for the follow-up session (Session 2), during which they completed the retention sequence block and the retention random block (50 trials each), followed by an intermanual transfer test, during which they completed the retention sequence block and the retention random block (50 trials each) with their untrained right hand.

Blinding efficacy for both the experimenter and the participant were assessed at the end of the first session, by asking the participant to complete a questionnaire which asked whether they thought they received (1) placebo or Madopar, and (2) sham, 1mA, 2mA, or 4mA stimulation. The experimenter also completed questionnaires assessing whether they thought the participant received (1) placebo or Madopar, and (2) sham or true stimulation.

### 2.5. Statistical Analyses

Due to poor performance (accuracy less than 60% during training), two subjects from the levodopa 4mA group were removed, and 1 subject from the levodopa 2mA group was removed. Reaction times from correct keypresses were used to estimate ***sequence-learning*** (also subsequently referred to as learning). We accounted for individual differences in reaction times by normalising reaction times to repeating sequences relative to reaction times in response to random sequences at baseline, prior to stimulation (i.e., from the baseline). Specifically, sequence learning was calculated by subtracting reaction times mean-averaged from sequence bins from reaction times mean-averaged from the baseline random bins (i.e., baseline mean random reaction time), divided by baseline mean random reaction time. This is expressed as follows: [(mean random reaction time – mean repeat reaction time)/mean random reaction time] (Perez, Wise, Willingham, & Cohen, 2007). Higher values indicate greater sequence-specific improvement. Sequence learning was quantified for baseline, acquisition, end of acquisition, retention, and intermanual transfer. We also quantified participants chronotypes via scores from the Morningness–Eveningness Questionnaire, and time of day (AM versus PM). These have not been included in the analyses as we met our stopping criteria with a sample we felt was too small to include covariates.

Bayesian ANOVAs and t-tests were run using JAMOVI, version 2.3.28.0. Specifically, for the training session, we ran Phase (p1, p2, p3, p4) x Bin (1….7) ANOVAs for sequence specific learning, calculated for each of the 28 training bins. For the no-drug, no-stimulation follow-up session, we ran Phase (p1, p2) x Bin (1…5) ANOVAs for sequence specific learning for the retention block and for the transfer block.

Bayes Factors (BF) values of 1–3 were interpreted as anecdotal, BF of 3–10 as moderate, and BF > 10 as strong evidence for the test hypothesis/variable inclusion (BF_10_ or BFincl), or for the null hypothesis/variable exclusion (BF_01_ or BFexcl). BF values ∼1 were interpreted as providing no evidential value. Statistical analyses were conducted using JAMOVI version 2.3.18.0.

## 3. Results

### 3.1. Baseline

Sequence specific learning did not differ across stimulation conditions prior to the start of the training phase and application of stimulation (main effect of Intensity, BF_excl_ = 15.244, Intensity x Drug interaction, BF_excl_ = 6.807). We found indeterminate evidence for the main effect of drug (BF_excl_ = 0.673).

### 3.2. Sequence learning

Overall, training improved performance. Specifically, reaction times for the sequence trials reduced pre-to post-training (BF_10,u_ = 3.49 e+55, cohen’s d = 2.24, 95% CI [2.02, 2.51]), in contrast to random trials (BF_10, u_ = 0.145, cohen’s d = 0.09, 95% CI [-0.10 0.29]), as Phase (Pre-training, Post-training) x Sequence (Sequence, Random) ANOVA showed a Phase x Sequence interaction (BF_incl_ = 2.48E+30). These benefits were retained after the > 48 hour delay, as reaction times for the first sequence bin remained faster than baseline (BF_10,u_ = 2.70 e+42, cohen’s d = 1.90, 95% CI [1.68 2.14]). A summary of the overall reaction times results is shown in Figure 5. Training-related performance improvements in sequence specific learning was evident across each training phase (main effect of phase, BFincl = ∞) as well as across bins (main effect of bin, BFincl =3.52e+52).

**Figure 5.**
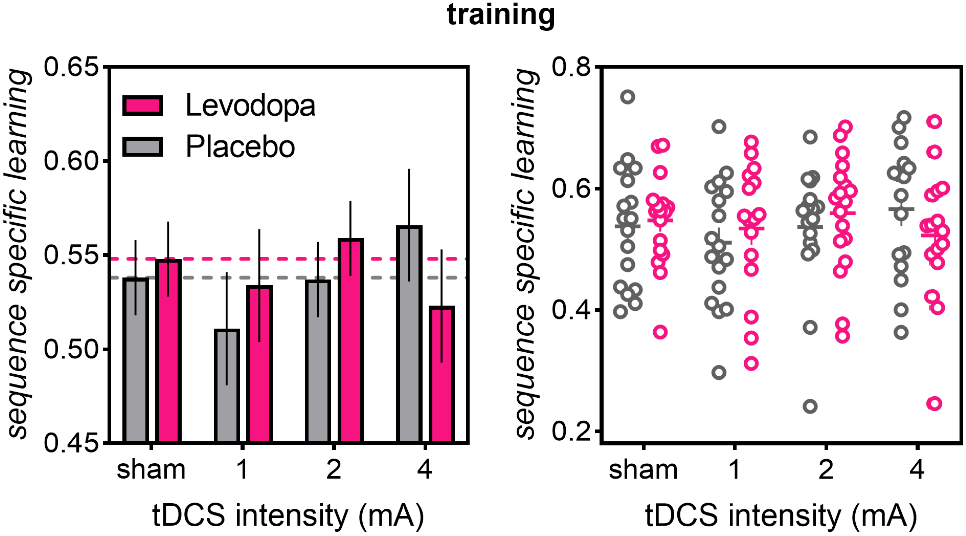
Sequence-learning during the training block. Higher values indicate greater sequence-learning. In the placebo group (gray bars), 4mA stimulation improved performance compared to all other groups, whereas 1mA stimulation resulted in worse performance than all other groups. In the levodopa group (pink bars), 4mA stimulation resulted in worse-than-sham performance, whereas 2mA stimulation resulted in better-than-sham performance. Dotted lines indicate performance for the sham stimulation group

### 3.2. Stimulation and levodopa modulated acquisition

Overall, in the absence of levodopa, 4mA tDCS improved – and 1mA tDCS impaired – acquisition of the sequence during training. The beneficial effect of 4mA tDCS was reversed with the administration of levodopa (see Figure 6). Indeed, acquisition across training phases was differentially altered by stimulation dose (Phase x Intensity interaction BF_incl_ = 173.690) as well as by the drug manipulation (Phase x Drug interaction BF _incl_ =18284.715). Follow-up Intensity (sham, 1mA, 2mA, 4mA) x Phase (Phase 1…4) x Bin (1…7) Bayesian ANOVAs run for the placebo group showed a Phase x Intensity interaction (BF_incl_ = 3.080), which was driven by improved acquisition for 4mA [(4mA vs sham: BF_10, U_ =12.131; 4mA vs 1mA: BF_10, U_ = 1.14E+07; 4mA vs 2mA: BF_10, U_ = 19.697), and impaired acquisition for 1mA (1mA vs sham: BF_10, U_ =8.9667, 1mA vs 2mA: BF_10, U_ = 5.1313, 1mA vs 4mA: BF10=1.14E+7). 2mA tDCS did not alter acquisition compared to sham (evidence for the null, BF_01, U_ = 13.583). The influence of levodopa on acquisition depended on stimulation intensity, (see Figure 6 left), as a follow-up Intensity x Phase x Bin ANOVA run for the levodopa group showed a Phase x Intensity interaction, BF_incl_ = 3.080. With levodopa, the 4mA group showed worse acquisition than the 2mA and sham groups (4mA vs sham: BF_10, U_ = 5.626; 4mA vs 2mA: BF_10, U_ = 615.994), and similar acquisition as 1mA (moderate evidence for the null: 4mA vs 1mA: BF_01, U_ = 6.096). 2mA tDCS improved acquisition compared to 1mA (BF_10, U_ = 7.676), but not compared to sham (moderate evidence for the null, BF_01, U_ = 4.050). The 1mA group showed similar learning as sham and 4mA, as shown by moderate evidence for the null (1mA vs sham: BF_01, U_ = 3.410; 1mA vs 4mA: BF01 _01, U_ = 6.09).

**Figure 6.**
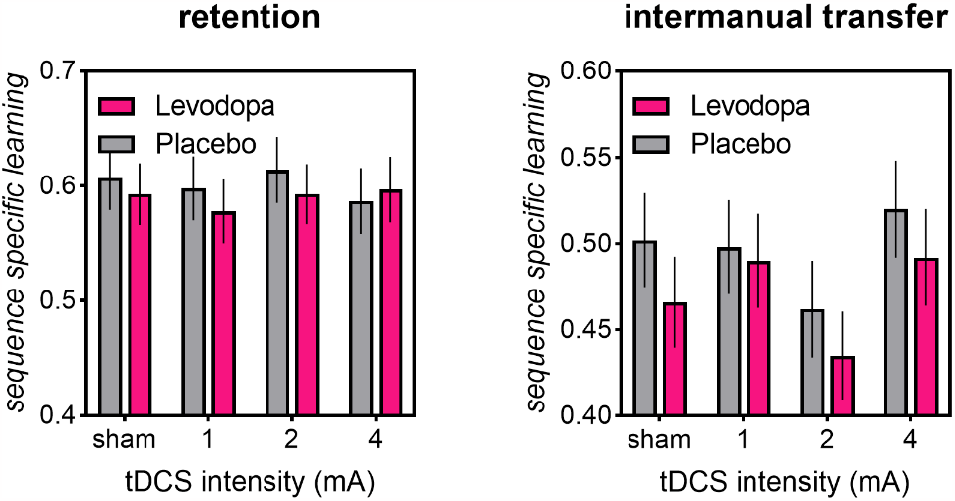
Sequence specific learning at follow-up, >48 hours after the training session. Whilst retention block performance was similar across stimulation and drug conditions, intermanual transfer assessed at follow-up showed poorest intermanual transfer for the 2mA condition for both the placebo group (gray bars) and the levodopa group (pink bars).

To quantify the effect of levodopa for each of the four intensities, we ran exploratory Drug x Phase x Bin ANOVAs (sham, 1mA, 2mA) separately for each intensity. Whilst the 4mA group showed impaired acquisition with levodopa than placebo [BF_10, U_ = 3827], the 1mA and the 2mA groups tended to show improved acquisition with levodopa than placebo, as shown by weak-moderate evidence for an effect of levodopa [1mA: BF_10, U_ = 2.42, 2mA: BF_10, U_ for = 3.61]. For sham tDCS, levodopa did not alter acquisition [moderate evidence for the null, BF_01, U_ = 6.61]

### 3.3. End of acquisition

Stimulation intensity and levodopa did not affect performance at the end of acquisition, as shown by moderate evidence for the null hypothesis (main effect of intensity: BF_excl_ = 8.849, main effect of Drug: BF_excl_ = 3.846, Drug x Hand interaction: BF_excl_ = 5.208, Drug x Hand x Intensity interaction BF_excl_ = 9.523).

### 3.4. Retention and intermanual transfer at follow-up

#### Retention

At follow-up, retention of sequence learning was evidenced in better-than-naive performance when comparing the 10 retention bins to the first 10 training bins (BF_10, U_ = 2.65e+155). Neither levodopa nor stimulation dose altered retention (main effect of Drug, BF_excl_ = 2.510, main effect of Intensity: BF_excl_ = 3.956, Drug x Intensity interaction: BF_excl_ = 3.460, Drug x Phase interaction, BF_excl_ = 9.509, Intensity x Phase interaction, BF_excl_ = 8.944). The absence of an effect of levodopa on tDCS effects at 48-hour follow-up is consistent with previous findings that levodopa augments the persistence of motor cortex tDCS effects on motor evoked potentials up to 48 hours (and not beyond 48 hours) after stimulation cessation (Kuo et al., 2008).

#### Intermanual transfer

Stimulation intensity at training altered intermanual transfer at follow-up. Whilst the main effect of intensity was inconclusive (BF_incl_ = 0.674), t-tests showed that across both the placebo and the levodopa group, 2mA tDCS resulted in poorest intermanual transfer compared to all other stimulation conditions (very strong evidence for the test hypothesis: 2mA vs sham: BF_10, U_ = 98.343, 2mA vs 1mA: BF_10, U_ =14176.472, 2mA vs 4mA, BF_10, U_ = 1.88e+7).

Levodopa at training did not alter intermanual transfer: we found inconclusive-to-moderate evidence for excluding the effect of the drug manipulation (main effect of Drug: BF_excl_ = 1.440, Drug x Intensity interaction, BF_excl_ = 8.264).

### 3.5. Blinding efficacy

Overall, blinding was successful for both experimenters and participants. Specifically, experimenters were not above chance at correctly guessing whether participants were assigned to the levodopa or placebo condition, or the verum versus active stimulation condition, as Bayesian binomial tests showed moderate evidence for the null hypothesis (drug condition: BF_01_ = 3.591, stimulation condition: BF_01_ = 5.445). Participants were not above chance at correctly guessing the drug condition (BF_01_ = 9.151) and not above chance at correctly guessing which one of the four stimulation conditions they were assigned to (BF_01_ = 8.669).

## 4. Discussion

In a pre-registered, double-blind study, using conservative Bayesian statistics, we found an intensity-dependent effect of anodal motor cortex tDCS on explicit motor learning that interacted with exogenous dopamine. Specifically, in the absence of levodopa, 1mA impaired, 2mA had no effect, and 4mA tDCS improved acquisition of motor sequence during training. Conversely, administration of levodopa reversed the beneficial effects of 4mA tDCS on acquisition. To the best of our knowledge, this is the first study to demonstrate a causal role of dopamine in the intensity-dependent effects of tDCS on behaviour.

The prominent benefit to learning in the highest intensity 4mA condition suggests that higher doses of injected current can be more potent in modulating function. This result is consistent with previous findings of large benefits of 4mA tDCS on sequence learning in neurotypical young adults (Hsu et al., 2023; Shinde et al., 2021). Similarly, positive dose-response relationships have been suggested from meta-analyses on clinical populations such as those who have had a stroke (Van Hoornweder et al., 2021) or major depression (Wang et al., 2021). An important caveat however is that the effect of stimulation intensity is likely task-dependent. For example, whilst linear effects of intensities have been shown for mind-wandering (Filmer, Griffin, & Dux, 2019a), non-linear effects of intensity have been shown for working memory training (Weller et al., 2020) and multitasking training (Ehrhardt et al., 2022). In a similar vein, the effect of stimulation intensity here depended on learning phase. Here, 4mA tDCS selectively improved whilst 1mA impaired acquisition without altering retention or intermanual transfer. In contrast, 2mA tDCS selectively impaired intermanual transfer without altering acquisition or retention. This parallels other work which found that different stimulation intensities differentially altered training and transfer (Ehrhardt et al., 2022).

We found that whilst levodopa resulted in a small benefit to acquisition of sequence learning when paired with lower tDCS intensities (1mA-2mA), levodopa reversed beneficial effects at the highest 4mA tDCS intensity. By showing that levodopa modulates effects of tDCS intensity on sequence learning, we demonstrate a causal role of dopamine in the way stimulation influences learning. Frontal cortex tDCS is known to induce midbrain dopamine release (Fonteneau et al., 2018; Fukai et al., 2019; Tanaka et al., 2013) and – at the dose used here – levodopa primarily increases midbrain dopamine (Carey, Pinheiro-Carrera, Dai, Tomaz, & Huston, 1995). A large body of evidence demonstrates an inverted U-shaped relationship between brain dopamine and cognition (for a review, see Cools & D’Esposito, 2011; Sawaguchi & Goldman-Rakic, 1991). Based on our findings showing that the effect of levodopa on sequence learning depends on the stimulation intensity used, it may be that dosage of tDCS differentially modulates dopamine release, which influences performance. Stimulation dose seems likely to modulate dopamine release: for example, moderate but not low or high intensities of repetitive transcranial magnetic stimulation resulted in dopamine release in the rat dorsolateral striatum (Kanno, Matsumoto, Togashi, Yoshioka, & Mano, 2004). According to this proposal, higher stimulation intensities might increase brain dopamine to optimal levels, but combining 4mA tDCS with levodopa results in excessive dopamine, impairing performance. This possibility remains to be shown experimentally.

Contrary to our predictions, we did not find evidence for an effect of levodopa on motor sequence learning in the absence of active tDCS. We predicted an effect of levodopa on sequence learning, based on previous studies linking dopamine genotypes and explicit motor sequence learning in older adults (Schuck et al., 2013), as well as effects of levodopa on explicit sequence learning in Parkinson’s disease (e.g.,Kwak, Muller, Bohnen, Dayalu, & Seidler, 2010). Profound differences in dopamine function between young and older adults (for a review, see Backman, Lindenberger, Li, & Nyberg, 2010) and between older adults and Parkinson’s disease patients (Collier, Kanaan, & Kordower, 2011) make it challenging to extrapolate expected findings across such populations. Our findings suggest that in young healthy adults, levodopa – at least with the dosage and the task used here – does not influence explicit motor sequence learning. It is noteworthy that recent work has similarly shown no effect of the dopamine D2 antagonist haloperidol on the learning of motor sequences in young adults: haloperidol only affected the speed of individual movements that comprised each sequence (Sporn & Galea, 2023). Future research is necessary to clarify if such findings will generalise to other populations with altered dopamine function.

Our findings have implications for the application of tDCS in populations with altered dopamine function, including older adults and Parkinson’s disease. In such populations, there is an increasing body of work demonstrating either null effects (Ghasemian-Shirvan et al., 2023; Horne et al., 2021; Tan, Filmer, & Dux, 2021) or unexpected negative effects of tDCS on behaviour (King et al., 2020; Puri et al., 2021; Simpson & Mak, 2022). For example, Ghasemian-Shirvan et al. (2023) found no effect of anodal 1mA, 2mA, 3mA motor cortex tDCS on motor sequence learning in elderly participants, even though a dose-dependent effect was prominent for motor evoked potentials. Stimulation applications in these populations may benefit from consideration of baseline dopamine function, potentially through pre-conditioning stimulation protocols, use of medications to modulate dopaminergic function (e.g., Perez, Morales-Quezada, & Fregni, 2014; Wang et al., 2014), and/or stimulation protocols designed to optimise outcomes specifically for these populations.

Previous imaging studies have shown that anodal motor cortex tDCS reduces concentrations of the inhibitory neurotransmitter Gamma-Aminobutyric Acid (GABA) in the contralateral and ipsilateral motor cortex (Bachtiar et al., 2018), and the extent of this decrease correlates with amount of improvement in sequence learning (Stagg, Bachtiar, & Johansen-Berg, 2011). tDCS to the frontal cortex is also known to result in a disinhibitory effect on the midbrain, by increasing striatal GABA levels (Bunai et al., 2021). Indeed the larger the striatal GABA increase, the larger the stimulation-induced dopamine release in the striatum (Bunai et al., 2021). Thus, interactions between the GABAergic and dopaminergic systems might give rise to the effect of tDCS on the cortex and behaviour. Emerging work has begun to combine manipulations of different neurochemicals to better understand how tDCS alters brain neurophysiology (Ghanavati, Salehinejad, De Melo, Nitsche, & Kuo, 2022). Future studies may combine measures/manipulations of GABA, dopamine, tDCS and stimulation dosage to disentangle and elucidate the combined contributions of these to behaviour.

In sum, we show non-linear dosage effects of tDCS on explicit motor sequence learning, and these interact with dopamine availability. This work represents a significant step forward in understanding how stimulation can modulate motor learning, giving mechanistic insights into neurochemical contributions to the efficacy of stimulation. Developments such as these will improve our ability to target stimulation to situations where it is more likely to be efficacious, and gives insight into how we may go about optimising applications in the future.

